# Intrinsic Cellular Persistent Firing Sustains Hippocampal Spatial Representations during Working Memory

**DOI:** 10.1101/2025.06.27.661493

**Authors:** Babak Saber Marouf, Antonio Reboreda, Frederik Theissen, Rahul Kaushik, Magdalena Sauvage, Alexander Dityatev, Motoharu Yoshida

**Affiliations:** Institute of Physiology, Medical Faculty, Otto-Von-Guericke University, 39120 Magdeburg, Germany.; Cognitive Neurophysiology, German Center for Neurodegenerative Diseases (DZNE), 39120 Magdeburg, Germany.; FAM Department, Leibniz Institute for Neurobiology (LIN), 39118 Magdeburg, Germany.; Molecular Neuroplasticity Group, German Center for Neurodegenerative Diseases (DZNE), 39120 Magdeburg, Germany.; Medical Faculty, Otto-von-Guericke University (OVGU), 39120 Magdeburg, Germany.; Center for Behavioral Brain Sciences (CBBS), 39106 Magdeburg, Germany.

## Abstract

Neurons are the fundamental units of cognitive processing. The prevailing view holds that neurons are input-output units that fire action potentials only upon receiving sufficient synaptic input. Here, we demonstrate that this may not be the sole operational mode of neurons *in vivo* during cognitive tasks, by showing that hippocampal place cells actively maintain firing and preserve spatial information through molecularly defined intrinsic mechanisms. Using *in vitro* and *in vivo* recordings coupled with molecular manipulation in mice, we reveal that hippocampal persistent firing during spatial working memory is supported by individual neurons via TRPC4 ion channels. Despite the general belief that persistent activity carries working memory content, we demonstrate that this firing maintains immediate spatial representations and is required for spatial working memory performance. These findings redefine neurons as active contributors to information retention, beyond their traditional role as passive input-output units, potentially reshaping our general understanding of computation in the brain.

## Introduction

Precise maintenance of internal representations in the brain is critical for executing cognitive tasks^1,2^. A potential cellular correlate for this process is persistent neural firing, in which information is maintained by continuous activation of a specific subset of cells^3,4^. Persistent firing has been observed across multiple brain regions, including the prefrontal cortex, hippocampus and sensory association areas, and is thought to support a wide range of cognitive functions such as working memory, decision-making, attention, spatial navigation, and goal-directed behavior^5–10,2^. In the rodent hippocampus, in addition to “time cells” that activate sequentially during relatively short delay intervals (∼10 s)^11,12^, a recent study has identified persistently firing cells that span the entire delay period (∼30 s) in the mouse hippocampal CA1, outlasting the time-cell sequence^13^.

Building on the foundational concepts of Lorente de Nó and Hebb, persistent activity has traditionally been attributed to reverberatory dynamics within recurrent synaptic networks^14,15^. This view aligns with the prevailing model of neurons as passive elements generating action potentials only when synaptic inputs exceed a threshold. However, these pure network models often exhibit instabilities such as noise-induced drift and runaway excitation^3^, and their performance depends on a degree of synaptic fine-tuning rarely observed in biological systems^4,16^. Consequently, such models often struggle to replicate the robust, long-lasting persistent firing observed in biological systems without incorporating additional stabilizing mechanisms^17^.

Neuromodulators such as acetylcholine, which is vital for working memory and other cognitive functions, can transform the way neurons respond to input signals^18^. *In vitro* studies have indicated that individual neurons can continue firing after a brief trigger under the influence of acetylcholine through an intrinsic cellular mechanism—a phenomenon referred to here as “intrinsic persistent firing”^19–24^. We and others have identified TRPC channels as key mediators of intrinsic persistent firing in CA1 pyramidal cells^20,25^. Theoretical studies have demonstrated potential roles for this cellular property in working memory and spatial coding^26,27^. Indeed, incorporating mechanisms of intrinsic persistent firing endows attractor network models with the high stability and robustness observed in *in vivo* data^28–30^. However, whether intrinsic persistent firing influences *in vivo* neural activity, sustains information, and supports cognitive function remains unclear because the effects of disrupting this property have not been examined in living animals^31,32^. As a consequence, current computational models—from classical attractor networks to the artificial neural networks underlying deep learning—largely rely on synaptic connectivity to sustain activity.

Beyond cellular mechanisms, the specific functional contribution of the hippocampus to spatial working memory remains a subject of intense debate. It has long been contested whether the hippocampus supports spatial working memory by maintaining task-specific content (working memory hypothesis) or by supplying an ongoing spatial representation (cognitive map theory)^33,34^. While human studies repeatedly indicate that hippocampal persistent firing encodes working memory content^8^, decoding analyses from the rodent hippocampus have shown an absence of working memory content (e.g. left vs right turns) during the delay period^13,35,36^. Thus, the information represented by hippocampal persistent firing during spatial working memory still remains unclear. Notably, both working memory maintenance and spatial representation have been modeled using similar attractor-based dynamics^37,38^, indicating that the maintenance of head direction or spatial information is associated with persistent firing^39,40^. However, whether the maintenance of spatial representations *in vivo* relies on network-level connectivity or individual cellular mechanisms remains to be determined.

Here, we test the contribution of intrinsically generated persistent firing to the maintenance of internal representations *in vivo* and to cognitive performance. We first identify the TRPC4 ion channel as a molecular mechanism underlying intrinsic persistent firing and subsequently examine neural activity in mice during a spatial working memory task under TRPC4 knockdown (KD). Persistent firing observed in control animals during the delay period of the task was markedly reduced in TRPC4 KD mice, indicating that intrinsic cellular mechanisms support persistent firing *in vivo*. Notably, the transient activity of cells before and after the delay period was not altered, highlighting the specific role of TRPC4 channels in sustained firing. Spatial working memory performance was significantly impaired, providing the first evidence for a functional role of hippocampal persistent firing in this task. Using support vector machine (SVM) decoding and spatial information measurements, we reveal that hippocampal persistent firing reflects the sustained activity of place cells with high spatial information when animals remain at fixed locations (the start and goal areas) for extended periods. Furthermore, using Bayesian decoding and SVM-based classification, we demonstrate that spatial representation errors increased rapidly and that left-right goal prediction deteriorated upon entry into the goal areas in TRPC4 KD mice, indicating impaired maintenance of spatial representations. Notably, the strength of persistent firing at the goal locations correlated with task performance. These data suggest that hippocampal persistent activity supports spatial working memory by maintaining instantaneous spatial representations. These results position individual neurons as active units of information retention, beyond their traditional role as passive input-output units, and offer a revised framework for understanding memory and neural computation.

## Results

### TRPC4 KD suppresses intrinsic persistent firing *in vitro*

We developed an shRNA TRPC4 channel knockdown (TRPC4 KD) virus and a control virus with a scrambled sequence (Scrambled) to test the role of intrinsic persistent firing *in vivo* (see Methods; Fig. 1A)^41^. The mice received bilateral injections of either virus into the CA1 region (Fig. 1B). First, a loss of intrinsic persistent firing was confirmed using *in vitro* patch-clamp recordings of hippocampal CA1 cells in the presence of the cholinergic receptor agonist carbachol^20,25^ (10 pM; Fig. 1C). Percentages of cells responded with persistent firing, the frequency of persistent firing and membrane depolarization were significantly lower in the TRPC4 KD group than in the two control groups (Fig. 1D-F). Basic properties of TRPC4 KD cells, such as the resting membrane potential, input resistance, adaptation index, and intrinsic excitability, did not significantly differ from those in the control groups (except for intrinsic excitability measured with 100 pA current injection, Kruskal-Wallis test, p = 0.0193, Chi-sq = 7.89, df = 2; Table 1). In summary, TRPC4 KD significantly decreased the ability of individual cells to support persistent firing while preserving basic cellular properties.

**Figure 1.**
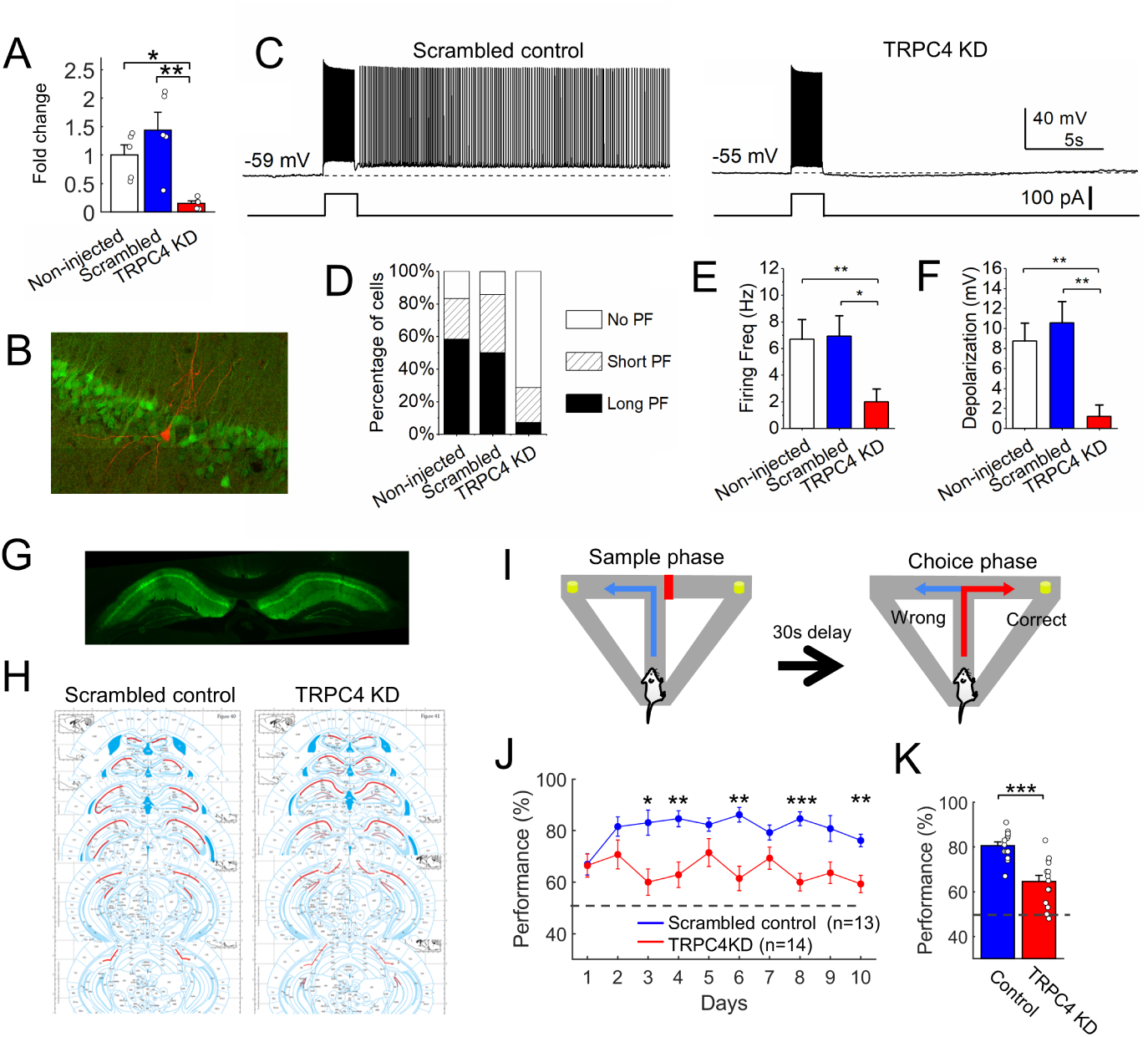
Hippocampal TRPC4 KD significantly reduces intrinsic persistent firing and impairs the delayed non-match-to-sample spatial working memory task (A) Reduced TRPC4 mRNA expression by the TRPC4 KD virus confirmed by RTq-PCR (one-way ANOVA, p < 0.01; Tukey post-hoc test corrected for multiple comparison, No virus vs Scrambled: p = 0.33, No virus vs TRPC4 KD: p < 0.05, Scrambled vs TRPC4 KD: p < 0.01; n = 5 in all groups). (B) An example hippocampal slice with GFP-expressing cells due to TRPC4 KD virus injection and a recorded neuron (red). (C) Persistent firing observed in a CA1 pyramidal cell infected with the scramble control virus (upper trace) and an absence of persistent firing in a cell infected with the TRPC4 KD virus (lower trace). The bottom trace indicates injected current. The red part of this trace corresponds to the 10 s interval when firing rate and depolarization were measured in E and F. (D) Percentages of cells responded with persistent firing (n = 12, 14 and 14, respectively; No PF: no persistent firing. Short PF: persistent firing which did not last for 30 s. Long PF: persistent firing lasted longer than 30 s). Fisher’s exact test, Non-injected vs Scrambled: p = 1; Non-injected vs TRPC4 KD: p = 0.0079; Scrambled vs TRPC4 KD: p = 0.0063. (E and F) TRPC4 KD significantly decreased frequency of persistent firing and membrane depolarization measured during the 10 s period after the triggering stimulus (Frequency: Kruskal Wallis test, p = 0.0023, Chi-sq = 12.18, df = 2; Tukey post-hoc test, Non-injected vs Scrambled: p = 0.99, Non-injected vs TRPC4 KD: p = 0.0077, Scrambled vs TRPC4 KD: p = 0.0074; Depolarization: Kruskal-Wallis test, p = 0.0116, Chi-sq = 8.92, df = 2; Tukey post-hoc test, Non-injected vs Scrambled: p = 0.89, Non-injected vs TRPC4 KD: p = 0.017, Scrambled vs TRPC4 KD: p = 0.047; n = 12, 14 and 14, respectively). KD, knockdown; ANOVA, analysis of variance. (G) An example of hippocampal GFP expression after TRPC4 KD virus injection. (H) Summary of virus expression along the rostro-caudal axis from all mice. Red lines indicate the maximum extent of virus spread in each rostro-caudal plane. (I) The delayed non-match-to-sample spatial working memory task in a T-maze. (J) Working memory performance in scrambled control and TRPC4 KD group animals over ten days. Two-way ANOVA suggested significant main effects of the animal group (p < 0.0001). T-tests corrected for multiple comparisons suggested significant differences on days 3 (p = 0.020), 4 (p = 0.0086), 6 (p = 0.0019), 8 (p < 0.001), and 10 (p = 0.0040). (K) Averaged working memory performances of TRPC4 KD mice were significantly lower than the scrambled control group (T-test, p < 0.001; n = 13 and 14, respectively). KD, knockdown; ANOVA, analysis of variance.

**Table 1.** Comparisons of passive membrane properties.

|  | RMP (mV) | Resistance<br>(MΩ) | Adaptation<br>Index | Intrinsic Excitability (Hz) |  |  |  |
| --- | --- | --- | --- | --- | --- | --- | --- |
|  |  |  |  | 50 pA | 100 pA | 150 pA | 200 pA |
| Non-injected | $-64.19 \pm 0.18$ | $163.31 \pm 4.82$ | $0.43 \pm 0.02$ | $5.09 \pm 0.48$ | $20.45 \pm 0.69$ | $29.55 \pm 0.71$ | $39 \pm 0.54$ |
| Scrambled | $-64.20 \pm 0.21$ | $180.66 \pm 2.72$ | $0.53 \pm 0.02$ | $2.5 \pm 0.35$ | $13.71 \pm 0.56$ | $25.21 \pm 0.64$ | $33.21 \pm 0.82$ |
| TRPC4 KD | $-63.79 \pm 0.25$ | $173.85 \pm 2.62$ | $0.67 \pm 0.03$ | $7.86 \pm 0.49$ | *$22.71 \pm 0.59$ | $30.5 \pm 0.70$ | $36.57 \pm 0.65$ |

RMP: resting membrane potential, Resistance: input resistance, Adaptation index: firing rate adaptation index.

### TRPC4 KD impairs spatial working memory

Second, the effect of TRPC4 KD on spatial working memory was assessed by a delayed non-match-to-sample paradigm using an automated T-maze (Fig. 1G-I; Supplemental Fig. 1). The scrambled control group mice consistently performed approximately 80% correctly from day two. On the other hand, the performance of the TRPC4 KD group remained approximately 60% correct over the ten days of testing (Fig. 1J). Group comparisons demonstrated that hippocampal TRPC4 KD significantly impaired working memory performance compared to scrambled control mice (Fig. 1K). These results indicate that TRPC4 channels in CA1 neurons are necessary for spatial working memory.

### TRPC4 KD suppresses *in vivo* persistent firing

Classical theory and evidence suggest that persistent firing supports memory maintenance^8,33,42^. We tested whether TRPC4 KD affected persistent firing in the T-maze using *in vivo* recordings. We observed cells with increasing firing rates during the delay (maintenance) period (0-30 s; orange line, Fig. 2A). To quantify such increased firing, we calculated the z-scored firing rate of individual cells around the delay period using the pre-delay period as the baseline in cells from the control and TRPC4 KD groups (Fig. 2B, see Methods). The cells were sorted based on their mean activity during the delay period (0-25 s; orange line at the top of Fig. 2B). While a substantial portion of cells at the top of the figure showed increased activity during the delay in the control group (top ∼10%; Fig. 2B, left), such an increase was less prominent in the TRPC4 KD group (Fig. 2B, right). The activity of the top 10% cells was significantly lower in the TRPC4 KD group (top panels in Fig. 2C and D). Similarly, suppression of the bottom 10% cells in the TRPC4 KD group was significantly weaker (bottom panels in Fig. 2C and D), indicating that both positive and negative modulations were compromised in the KD group. Accordingly, TRPC4 KD significantly reduced the number of delay-activated (Act) and delay-suppressed (Sup) cells and increased the number of non-modulated (NM) cells (Fig. 2E).

**Figure 2.**
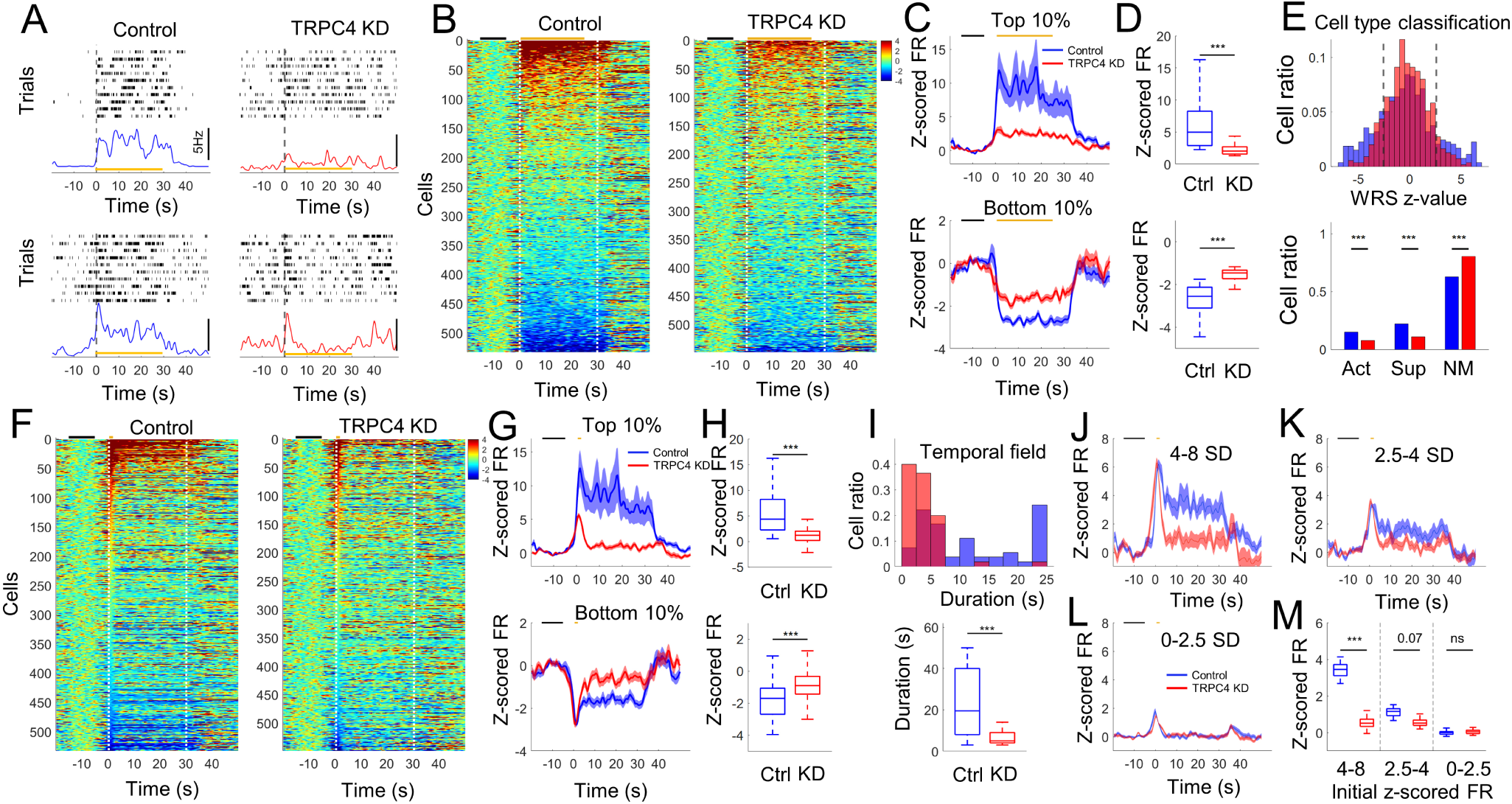
TRPC4 KD suppresses in vivo persistent activity (A) Example firing pattern (raster plot) and averaged firing rate (lower panel) of two cells from the control and TRPC4 KD groups around the delay period of the spatial working memory task. (B) Z-scored firing rate of all cells sorted by the delay period activity. Vertical dashed lines indicate the onset and offset of the delay period. Black and orange horizontal lines at the top of each panel indicate the baseline period and period for cell sorting, respectively. (C) Averaged z-scored firing rate (FR) of the top (upper panel) and bottom (lower panel) 10% cells in B. (D) Comparison of z-scored firing rate of the top (upper panel) and bottom (lower panel) 10 % cells in B averaged over the delay period (0-25 s) (WRS test, p < 0.001 and p < 0.001 for upper and lower panels respectively). (E) Classification of activated (Act), suppressed (Sup), and non-modulated (NM) cells using the Wilcoxon rank sum (WRS) test. The upper panel shows the distribution of WRS z-values, and the lower panel shows the ratio of cells in each category (Chi-square test, p<0.001 for all categories). (F) Z-scored firing rate of all cells sorted by the activity at the initial part of the delay period (0-2 s). Black and orange horizontal lines at the top of each panel indicate the baseline period and period for cell sorting, respectively. (G) Averaged z-scored firing rate of top (upper panel) and bottom (lower panel) 10% cells in F. (H) Comparison of z-scored firing rate of the top (upper panel) and bottom (lower panel) 10% cells in F averaged over the delay period (0-25 s) (WRS test, p < 0.001 and p < 0.001 for upper and lower panels respectively). (I) Distribution (upper panel) and comparison (lower panel) of temporal field duration among the top 10% cells (WRS test, p < 0.001). (J-L) Averaged z-scored firing rate of cells that showed 4-8 (J), 2.5-4 (K), and 0-2.5 (L) SD peak z-scored firing rate at the initial part of the delay (0-2 s). (M) Comparison of activity during the delay (4-25 s) period in cells that showed different levels of initial activation corresponding to J to L (WRS test, p < 0.001, p = 0.07, and p = 0.58 for J, K, and L, respectively). KD, knockdown

If intrinsic persistent firing was supporting *in vivo* persistent firing, TRPC4 KD should increase transiently active cells. To test this, we sorted the cells using only activity during the first two seconds of the delay period (0-2 s, Fig. 2F). The activity of initially active cells significantly decreased during the delay period in the KD group, whereas more cells in the control group maintained their activity until the end of the delay period (Fig. 2F and G). The activity measured during the entire delay period (0-25 s) was significantly lower in the KD group (Fig. 2H). We measured the temporal field duration to quantify the duration of continuous elevation of the firing rate (Fig. 2I; see Methods). While the control group showed a wide distribution of temporal field duration up to the full delay duration, the majority of the TRPC4 KD group cells showed a temporal field shorter than 7.5 s (Fig. 2I, upper panel). The mean temporal field duration was significantly shorter in the TRPC4 KD group (Fig. 2I, lower panel). These data indicated that TRPC4 KD increased transient firing *in vivo,* as expected from our *in vitro* results.

However, the initial activity of the top 10% cells was stronger in the control group (Fig. 2G, upper panel), which could have resulted in stronger persistent firing in this group. To test this possibility, we filtered cells based on their initial (0-2 s) activity levels (Fig. 2J-M). The control group cells with initial activity increases of 4-8 and 2.5-4 standard deviations (SD) maintained a greater elevated level of activity until the end of delay compared to KD group cells (Fig. 2J and K), whereas those with initial activity of 0-2.5 SD did not show elevated activity (Fig. 2L). The sustained activity measured at 4-25 s was smaller in the KD group with 4-8 and 2.5-4 initial SD (Fig. 2M), indicating that the different activities between the two groups are not simply due to the difference in initial activation. Thus, more KD cells showed transient activity during the delay period, supporting the contribution of TRPC4 channels to sustained activity *in vivo* in a similar manner as *in vitro*.

### Temporal sorting reveals reduced persistent firing in TRPC4 KD mice

We further characterized the effect of TRPC4 KD using temporal sorting^11,12^. For this analysis, we selected putative pyramidal cells active (firing rate > 2 Hz) in at least one bin during or near the delay period (see Methods, 196/533 control cells and 235/549 TRPC4 KD cells) and sorted them according to their activity peak time, as in previous reports^11^ (Fig. 3A). We found an overrepresentation of the initial part of the delay period (0-3 s) in both the control and TRPC4 KD group (Fig. 3B). However, TRPC4 KD did not significantly alter peak time distribution.

**Figure. 3.**
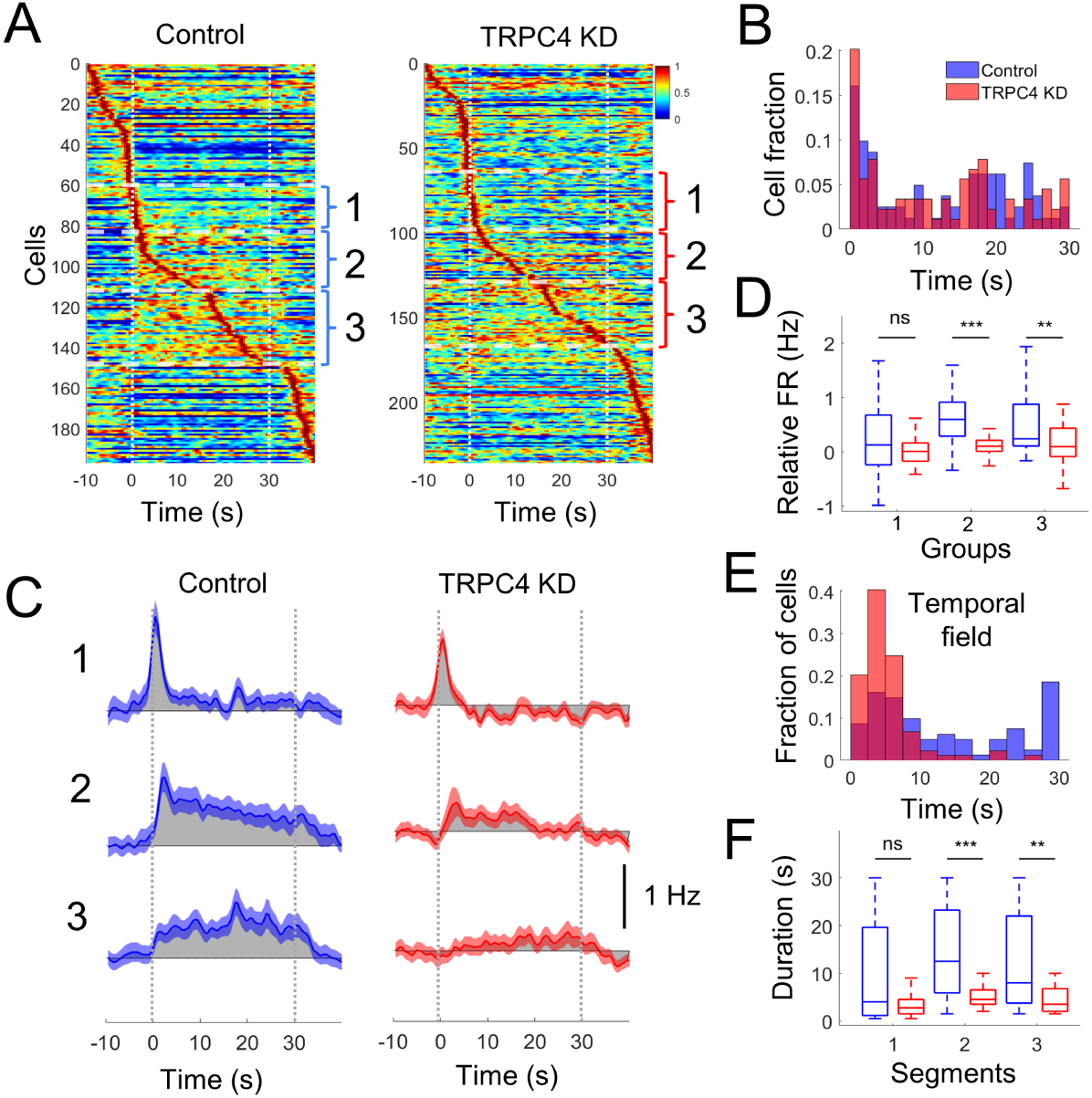
Peak time sorting also shows reduced persistent activity in TRPC4 KD mice (A) Normalized firing rate of cells sorted by their peak times. Vertical dashed lines indicate the onset and offset of the delay period. Horizontal dashed lines and numbers (1-3) on the right of each panel indicate divisions of cells by their peak times (group 1: 0-2.5 s, group 2: 2.5-16.5 s, and group 3: 16.5-30 s). (B) Distribution of peak time of all cells within the delay period. Peak time distribution is not significantly affected by TRPC4 KD (WRS test, p = 0.74). (C) Averaged relative firing rate (compared to baseline firing rate) of cells belonging to each peak time segment shown in A. Gray shades indicate a difference from the baseline firing rate. Note, a prominent, long-lasting elevated firing rate in control cells in groups 2 and 3. (D) Relative firing rate (FR) measured during the delay (0 to 30 s) compared to the baseline period in cells in each peak time segment. (E) Distribution of temporal field duration of all cells peaked during the delay period (group 1-3) (WRS test, p < 0.001). (F) Comparison of the duration of the field in each group (WRS test, p = 0.32, p < 0.001, and p = 0.006 for groups 1, 2, and 3, respectively). KD, knockdown

To evaluate potential effects of TRPC4 KD on cells with different peak times, cells were separated into three groups based on their peak times during the delay (Fig. 3A). While cells that peaked at 0-2.5 s (group 1) showed transient peak activity, the other two groups that peaked later in the delay (2.5-16.5 s: group 2, 16.5-30 s: group 3) exhibited, on average, an elevated firing rate throughout the delay period in the control group (Fig. 3C left). While TRPC4 KD did not affect the transient activation of cells at the initial part of the delay (group 1), it reduced long-lasting elevated firing of group 2 and 3 cells (Fig. 3C, right). Accordingly, the firing rate averaged over the delay period was significantly lower in groups 2 and 3 of the KD group but not in group 1 (Fig. 3D). Combining all cells that peaked during the delay, the KD group had more cells with shorter fields (See Methods; Fig. 3E). The duration of the field from the KD group cells was significantly shorter in groups 2 and 3 (Fig. 3F). The specific effect of TRPC4 KD on cells with longer temporal fields, further support the idea that the ability of single neurons to sustain firing is necessary specifically for persistent firing *in vivo*.

### Activity outside the delay period is intact

To further assess whether the effect of TRPC4 KD was specific to the delay period, we compared the activity of cells at different time points, ranging from 4 s before delay onset to 3 s after the delay (Fig. 4). First, cells were sorted using the z-scored activity of each cell at 3 to 4 s before the start of the delay period (−4 to −3 s; Fig. 4A), and the average activity of the top 10% cells from the control and KD groups was compared (Fig. 4D most left). These cells were only temporarily active and did not display persistent firing in either group, indicating that persistent firing was not ubiquitous. Neither activity of the top 10 % cells during this 1 s window (−4 s in Fig. 4E) nor during the entire delay period (−4 s in Fig. 4F) differed between the control and KD groups, indicating that TRPC4 KD did not have a strong effect before the delay. Similar results were obtained 2 s before delay onset (−2 to −1 s; Fig. 4D, second from left; Time −2 s in Fig. 4E and F). In contrast, control group cells activated during the delay (0-1, 10-11 and 20-21 s) showed stronger activity than KD cells, both during these 1s windows and during the entire delay (Fig. 4B; Fig. 4D middle three panels; Time 0, 10, and 20 s in Fig. 4E and F). Finally, when active cells were detected after the delay period (1-2 and 3-4 s later; Fig. 4C; Fig. 4D, two rightmost panels), the activity of these cells was transient and similar between the two groups (Time 1 and 3 s in Fig. 4E and F). This suggests that persistent firing does not occur before or after the delay period, and that TRPC4 KD specifically affects the activity of neurons during the delay period, as opposed to worsening the signal-to-noise ratio throughout the task.

**Figure 4.**
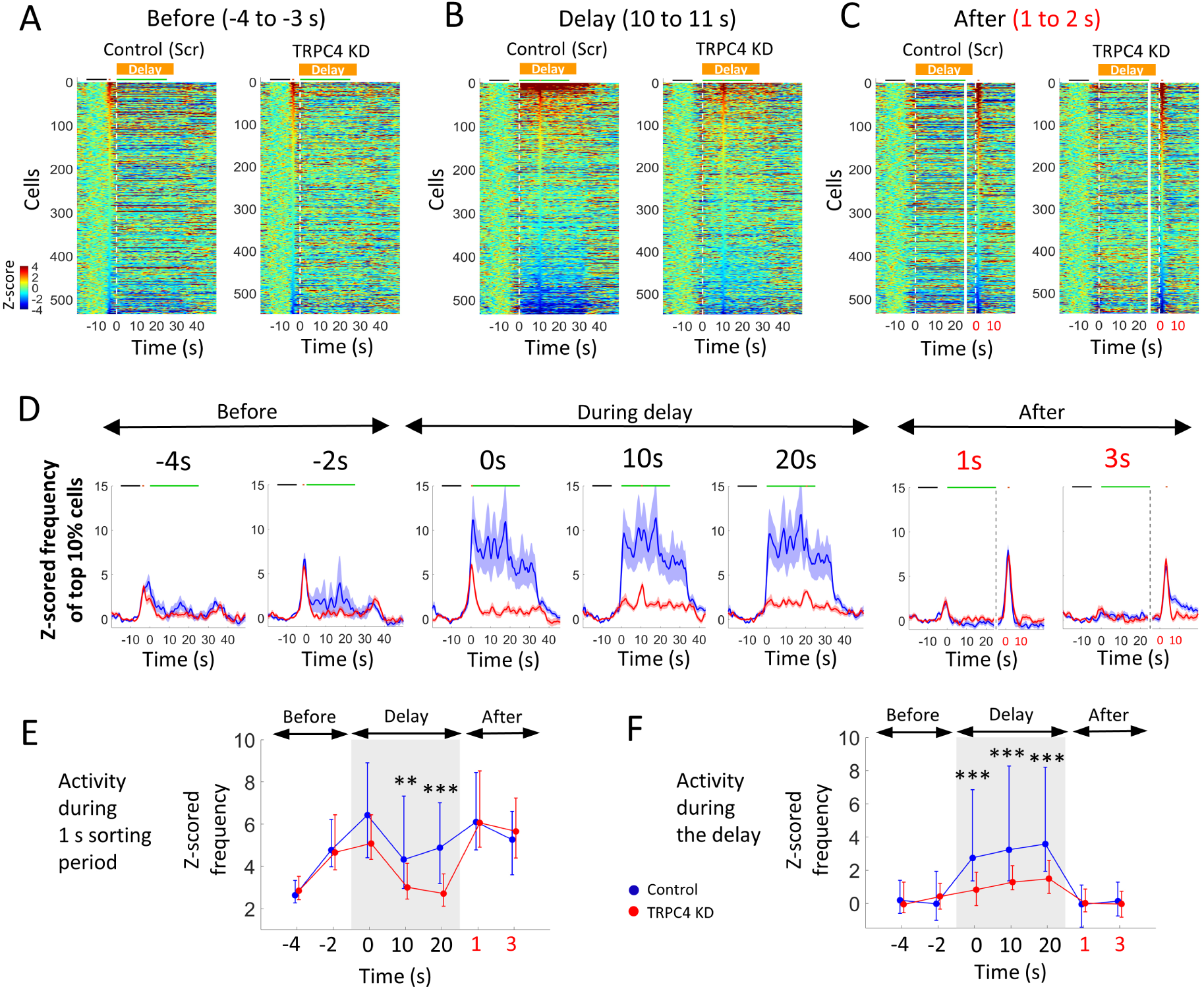
Activity outside the delay period is not strongly affected by TRPC4 KD (A-C) Z-scored firing rate of all cells sorted by their activity during the (A) 4 to 3 s before delay, (B) 10 to 11 s from the start of delay, and (C) 1 to 2 s after the delay offset. Vertical dashed lines indicate the onset and offset of the delay period. Black, orange, and green horizontal lines at the top of each panel indicate the baseline period, period for cell sorting, and period for delay activity measurement, respectively. In C, the white solid vertical line indicates the separation of the color plot into two parts that are aligned to the delay onset (left of the line) and offset (right of the line). (D) Averaged z-scored firing rate of the top 10% cells at different time points. Note that the cells chosen for each panel are therefore different. The vertical dotted lines in the two last panels correspond to the time when data alignment changed as in C. (E) Comparison of z-scored firing rate of the top 10 % cells measured specifically during the one second-long sorting period (corresponding to the orange horizontal lines in D; Ranksum test with Bonferroni correction, p = 0.31, 1.45, 0.29, 0.003, < 0.001, 1.45, 0.28, respectively). (F) Comparison of z-scored firing rate of the top 10 % cells measured specifically during the delay period (corresponding to the green horizontal lines in D; Ranksum test with Bonferroni correction, p = 0.88,1.46, < 0.001, < 0.001, < 0.001, 1.47, 1.10, respectively). Data points show the median, and error bars indicate the upper and lower quartiles. KD, knockdown

### Activity of putative pyramidal cells and interneurons

We next investigated whether the difference in the delay period activity between control and TRPC4 KD mice was cell type-dependent. We separated recorded neurons into putative pyramidal neurons and interneurons based on their average firing rates and spike durations (see Methods). Putative pyramidal cells in the control group showed strong activity during the delay, with TRPC4 KD suppressing this activity (Supplemental Fig. 2A-D). On the other hand, the activity of the top 10% putative interneurons during the delay period did not differ between the control and TRPC4 KD groups (Supplemental Fig. 2E-H). This suggests that putative interneurons are not the main source of the observed TRPC4 KD-dependent changes.

### Information coded by persistent firing

Next, we investigated the nature of information coded by cells with strong persistent firing during the delay period. The hippocampus may support spatial working memory in at least two ways. First, persistent firing may sustain working memory content (e.g., turn direction in the sample trial) during the delay period^33,42^. Second, the hippocampus may support spatial working memory tasks by providing information of the current spatial location^34,43^. However, whether the long-lasting persistent firing codes for the turn direction or current spatial information remains unclear.

We first compared the activity of cells in the control group during the delay period with that during the inter-trial interval (ITI; Fig. 5A). Since the animals were in the same start area in both the delay period and ITI, the different activities in these phases may suggest the coding of task-relevant information. Cells were sorted according to their activity during the delay period (Fig. 5B). Control cells highly active during the delay period were strongly active during ITI as well, without significant difference, as shown in Fig. 5B and C. The activity of individual cells during the delay period was similar to that during ITI, as shown in Fig. 5D.

**Figure 5.**
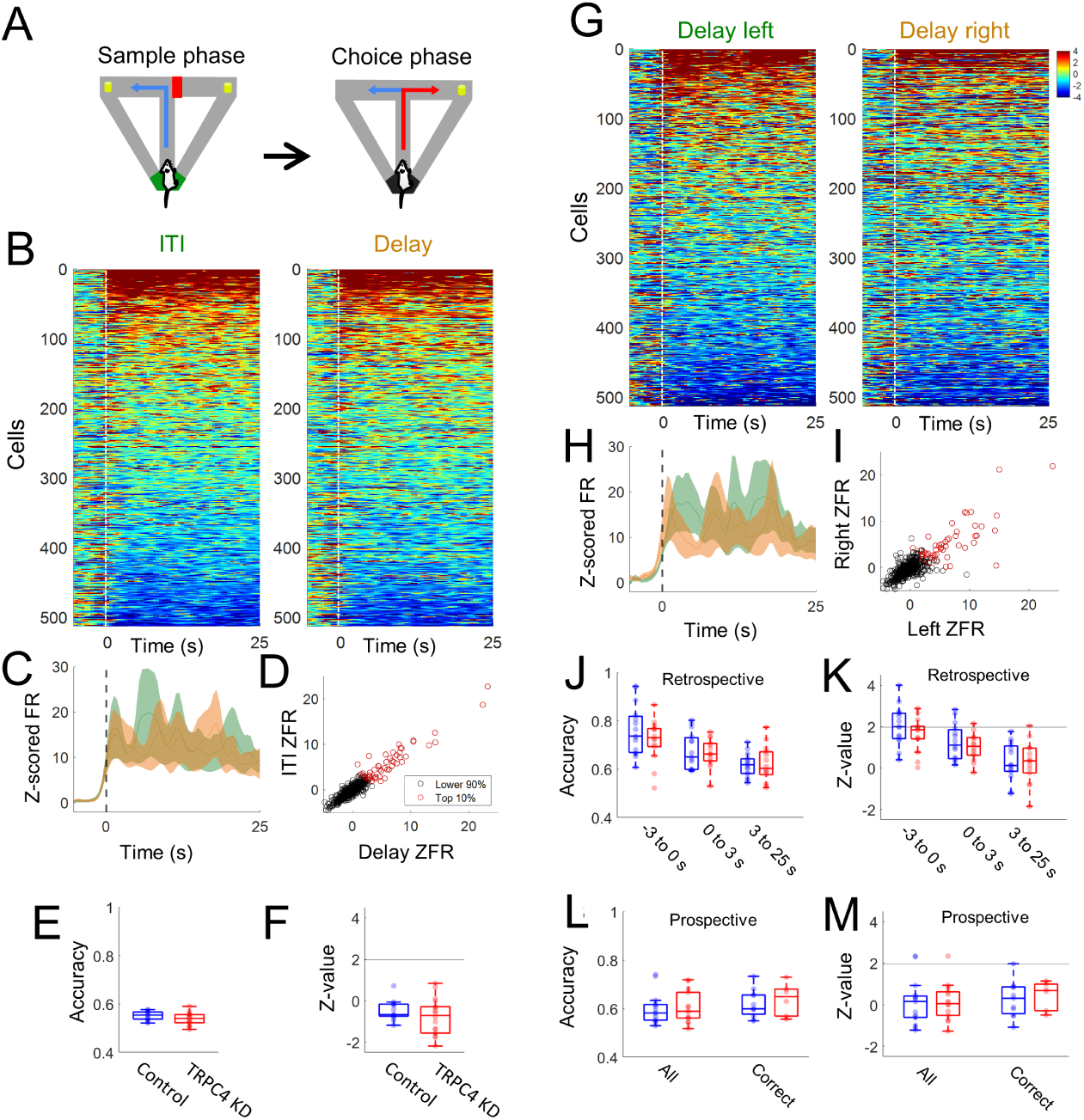
Persistently firing cells do not strongly code task phases and direction of turn (A) Inter-trial-interval (ITI, green) in the sample phase and delay period (orange) in the choice phase. Animals were in the same start box in the ITI and delay period. (B) Z-scored firing rate of all cells sorted by the averaged activity during the ITI (0-25 s) in both ITI and delay. Vertical dashed lines indicate the onset of ITI or the delay period. Baseline and sorting period were the same as in Fig. 2B. (C) Averaged activity of the top 10% cells in B from ITI (green) and delay period (orange; WRS test, p = 0.40). (D) Comparison of activity during ITI and the delay period from all cells. Each circle corresponds to a cell. The top 10% cells (based on common sorting) are indicated by red circles. (E-F) Accuracy (E) and z-value (F) from the SVM prediction of ITI vs delay phase based on the activity of all cells (WRS test, p = 0.61 and p = 0.59 for E and F, respectively). The horizontal line in F indicates a 5% significance level. (G) Z-scored firing rate of all cells near the delay period, sorted by the average activity of the left trials. Here, the left and right trials are defined by the direction of the turn at the sample phase (retrospective coding). (H) Averaged activity of the top 10 % cells in G from the left (green) and right (orange) trials. (WRS test, p = 0.0058). (I) Comparison of activity from left and right trials for all cells. Each circle corresponds to a cell. The top 10% cells (based on common sorting) are indicated by the red circles. (J-K) Accuracy (J) and z-value (K) from the SVM prediction of left vs right trials based on the activity of all cells at different segments before (−3 to 0) and during (0 to 3s, 3 to 25 s) the delay period (WRS test, p = 0.49, 0.91 and 1 in J; p = 0.20, 0.28 and 0.98 in K). (L-M) Accuracy (L) and z-value (M) from the SVM prediction of future left vs right trials based on the activity of all cells during (3 to 25 s) the delay period (prospective coding). This analysis was done using all (correct and wrong) and only correct trials (WRS test, p = 0.98 and p = 0.59, respectively, in L; WRS test, p = 0.78 and p = 0.52, respectively, in M). SVM, support vector machine

We further used a support vector machine (SVM) to test whether it could be trained to predict the delay and ITI phases based on the activity of all cells in each recording session (control: n = 16, TRPC4 KD: n = 18 sessions; see Methods for details; Fig. 5E and F). SVM could not differentiate the delay vs. ITI phases in any of the recordings (Z-values were lower than 1.96) from the control and TRPC KD groups. Furthermore, there were no differences between the control and KD groups. Together, these findings contradicted the idea that persistent firing codes for task-phase-relevant information.

Second, we asked whether the direction of the past or future turns was coded in persistent activity observed during the delay period. We sorted cells from the control group using the delay activity of the left trials (left turn on the sample trial; Fig. 5G, left) and used the same order for the delay activity of the right trials (Fig. 5G, right). A small but significant difference was observed when comparing the activity of the top 10% cells (Fig. 5H). The activity of individual cells in the left and right delay periods was generally similar but showed more differences than the comparison between the ITI and delay phases (Fig. 5I). While this difference between the left and right delay activity indicates the potential existence of coding of the turn direction, random noise may result in such differences.

Therefore, we further used an SVM to test whether the population activity codes for the left vs. right turn in each recording session (control, n = 15; TRPC4 KD, n = 14). SVM prediction accuracy exceeded 70% in most of the sessions (Fig. 5J) just before the onset of the delay period (−3 to 0 s), when the animal was still in the return arm of the T-maze, and this prediction accuracy was significantly higher than the chance level in approximately half (53.3%) of the sessions (Fig. 5K). However, prediction performance decreased quickly as the delay started (0-3 s; Fig. 5J and K), and almost none of the sessions reached a significant level in either the control or TRPC KD group after 3 s (3-25 s) in the delay period. No difference was observed between the SVM performance of the control and KD groups for any of the three time-segments.

We further asked whether the delay period activity codes for the future turn direction (prospective coding) rather than the past turn direction from the sample phase (retrospective coding). The SVM prediction for prospective coding was tested using the activity from the 3-25 s period during the delay of all or only correct trials (Fig. 5L and M). Most of the session did not reach significant levels both in the control and KD group and there was no difference between the control and KD. Together, these suggest that coding for the direction of turns during the delay period is weak. These agree with recent publications which pointed out that hippocampal delay period activity does not^35,44^ or only weakly^45^ codes for the direction of the past or future turns.

### Persistently active cells encode spatial information

Next, we tested the hypothesis that persistently active cells primarily code for spatial information. Computational models of hippocampal place cells suggest that spatial representation is maintained by the self-sustained activity of a subset of place cells^40^; therefore, persistent firing might be involved in the maintenance of spatial representation. We first sorted cells in order of the strength of persistent firing during the delay period as in the previous sections (Fig. 6A). We then calculated the spatial information of each cell using the entire duration of the recording, including the sample and choice phases from all 10 trials (Fig. 6B). Spatial information was particularly high in the control group cells which showed strong persistent firing (blue line, Fig. 6B). Spatial information of the top 10% cells in Fig. 6A was significantly lower in the KD group compared to the control group (Fig. 6C), whereas that of the bottom 10% cells was significantly higher in the KD group (Fig. 6D).

**Figure 6.**
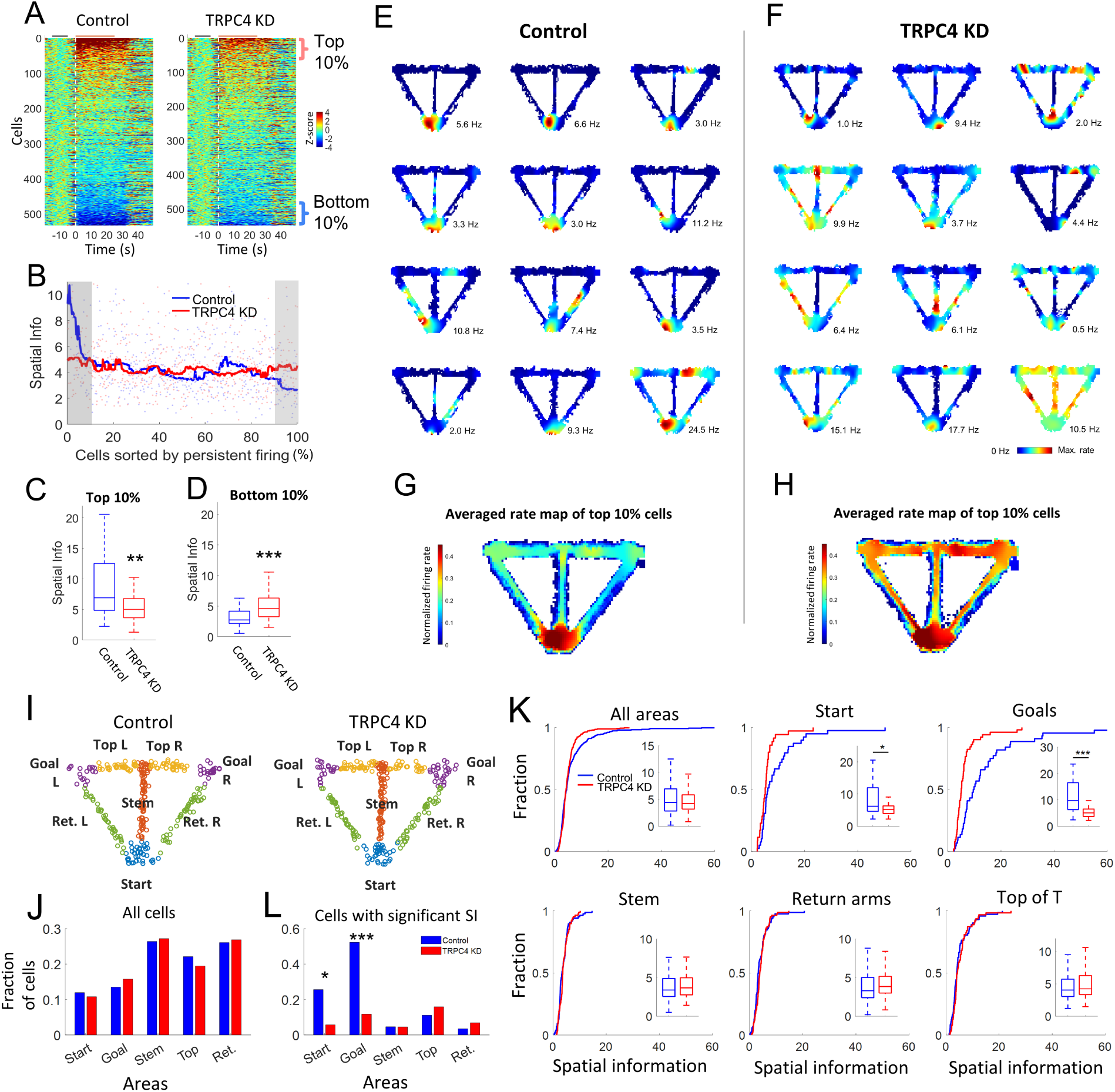
TRPC4 supports persistent activity of spatial cells when animals stay in a place for an extended duration (A) Z-scored firing rate of all cells sorted by the averaged activity during the delay period (0-25 s), as in Fig. 2B. Vertical dashed lines indicate the onset of the delay period. (B) Spatial information of all cells plotted in the order of the sorting in A. Solid lines indicate the moving average. Red and blue shaded areas indicate the top and bottom 10% cells. (C and D) Comparison of spatial information between the control and TRPC4 KD groups in the top (C; WRS test, p = 0.0024; n = 53) and bottom (D; WRS test, p < 0.001; n = 55) 10% cells. (E and F) Example spatial rate maps of the top 10% cells in control (E) and TRPC4 KD (F) groups. The maximum firing rate of each cell is indicated at the bottom right corner of each panel. (G and H) Averaged spatial rate maps of all top 10 % cells in the control (G) and TRPC4 KD (F) groups. (I) Distribution of peak firing location of all putative pyramidal cells on the T-maze. Each circle corresponds to a cell, and colors indicate different sub-areas. Top: top of the T-maze, Ret.: return arm, L: Left, R: Right. (J) Fractions of cells classified into each sub-area. (K) Distribution of spatial information of cells in all and each sub-area. Insets are comparisons of spatial information between the control and KD cells (WRS test, All areas: p = 0.56, Start: p < 0.019, Goals: p < 0.001, Stem: p = 0.41, Return arms: p = 0.19, and Top of T: 0.35). (L) Number of place cells with significant spatial information in each sub-area (Fisher’s exact test, p = 0.027, < 0.001, 1,0.45, and 0.49, respectively, from left to right). KD, knockdown

We visualized the spatial rate maps of the top 10% cells in control and TRPC4 KD groups (Fig. 6E and F). While the control group cells generally showed a clear field within the start area, fields of TRPC4 KD cells were less confined, often having multiple fields outside the start area (Fig. 6E and F). The averaged rate map from top 10% control cells showed a distinct firing at the start area, while that from the KD cells showed spatially less defined activity (Fig. 6G and H). This suggests that cells with strong persistent firing during the delay period are those with better-defined place fields in the start area and consequently have higher spatial information. This suggests that TRPC4 KD destabilized spatial representation by preventing persistent activity of CA1 place cells while the animal remained in the start area.

### Selective decrease of spatial information in start and goal areas

In the T-maze task, in addition to the start area, the animals stayed for an extended period in the goal areas where they consumed their rewards, whereas they run through other subareas of the maze (e.g., stem, return arm). If intrinsic persistent firing supports continuous place cell activity, TRPC4 KD might have affected spatial representation specifically in the start and goal areas without affecting that in other subareas. To test this, we first detected the location of firing fields within the T-maze for each putative pyramidal cell (see Methods). Next, we divided cells into different groups depending on the subareas of the T-maze where the location of field centers fell (Fig. 6I). The number of cells in each subarea was similar in the control and KD groups (Fig. 6J).

We then compared the spatial information of the cells in each subarea (Fig. 6K). Spatial information was slightly smaller in the TRPC4 KD group when all subareas were combined (all areas), without a significant difference. Consistent with previous observations, the spatial information of the cells in the start area was significantly lower in the KD group. Interestingly, a similar difference was observed in the goal area, where the animals stayed for an extended time. However, spatial information was not significantly different in other areas, such as the stem, return arms, and top of the T. Moreover, the number of place cells (cells significantly modulated by location) was higher in the control group than in the KD group only in the start and goal areas (Fig. 6L). These results suggest that TRPC4 KD selectively impairs the spatial representation when animals stay in a place, and place cells need to fire persistently. These suggest that TRPC4 KD specifically affects the maintenance of spatial representation when animal remained at the same place.

Additional analyses indicated that the compromised spatial representation observed in the start and goal areas in the TRPC4 KD animals is not associated with a lower running speed or weaker theta oscillations (Supplemental Fig. 3). These spatial analyses are in line with the time-domain analysis (Fig. 4), which revealed a specific effect of TRPC4 KD on cellular activity during, but not before or after, the delay.

### Gradual decrease of left and right goal representation

We further analyzed cellular activity and spatial representation in the goal areas (Fig. 7) since the encoding of left vs. right goals in the hippocampus during the sample phase may be crucial for task performance^43^. First, we tested whether persistent activity differed in the goal areas. We limited our analysis to trials in which animals spent at least 10 s continuously within the goal area (see Methods for details). Fig. 7A shows the z-scored firing rate of all cells where the time 0-10 s corresponds to the 10 s period when animals continuously stayed in the left goal area in the sample phases (indicated by the orange line on the top). Activity of the top 10% cells was significantly lower in the TRPC4 KD group, both in the left and right goals (Fig. 7B and C). These results indicated that TRPC4 KD reduced the continuous activity of cells in the goal area as in the start area.

**Figure 7.**
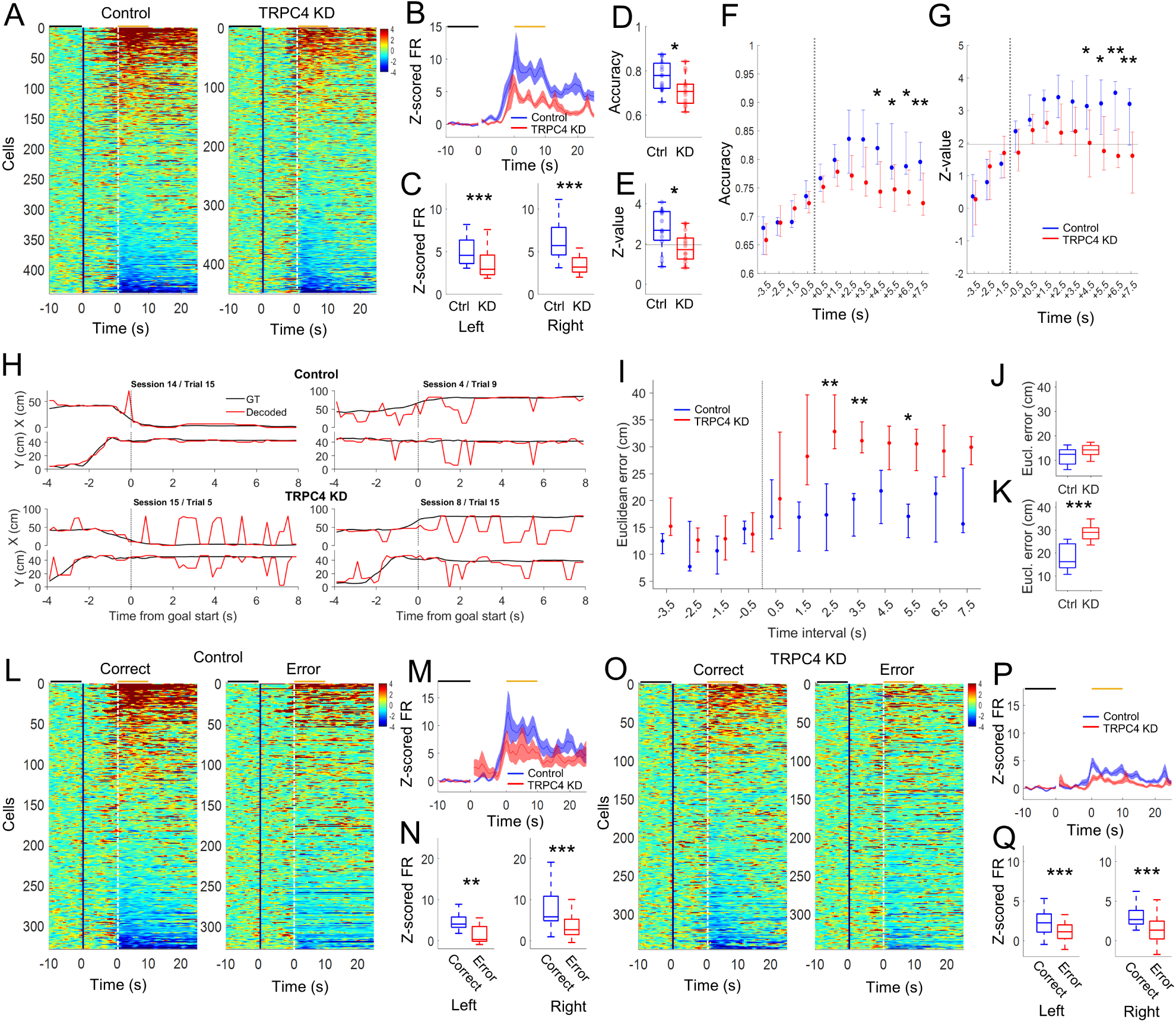
Gradual decrease of spatial representation in the goal areas may have resulted in the poor task performance of TRPC4 KD mice (A) Z-scored firing rate of cells sorted by their activity during the 10 s goal period when animals continuously stayed in the left goal area. The black vertical gap separates the first 10 s, which is referenced to the end of the ITI period, and the rest, which is referenced to the start of the goal entry. The black and orange horizontal lines on the top indicate the baseline and 10 s goal period, respectively. (B) Activity of the top 10% cells in A. (C) Comparisons of activity of the top 10% cells during the 10 s goal period (0-10 s) at the left and right goals (WRS test, p < 0.001 and p < 0.001, respectively). (D-E) Accuracy (D) and z-value (E) from the SVM prediction of left vs right goals based on the activity of all cells in individual sessions during the 10 s goal period (WRS test, p = 0.019 and 0.017 in D and E, respectively). The horizontal line in E indicates a 5% significance level. (F-G) Accuracy (F) and z-value (G) from the SVM prediction of left vs right goals at different time periods before and during the 10 s goal period. The vertical dotted lines indicate goal entry, and the horizontal line in G indicates 5% significance level. (H) Example real (GT) and decoded X and Y positions of mice from 4 s before to 8 s after goal entry. Dashed lines indicate time of goal entry. (I) Decoding error measured as distance between true and estimated positions at different time periods before and during the goal period. (J and K) Comparison of error distance between the control and the KD groups before (−4 to 0 s) and after (0 to 8 s) goal entry (WRS test, p = 0.19 and 0.001, respectively), (L) Z-scored firing rate of cells from the control group animals sorted by their combined sort order (see text) during the 10 s goal period in the left goal area from the correct (left panel) and error (right panel) trials. (M) Activity of the top 10% cells in L. (N) Comparisons of activity of the top 10% cells during the 10 s goal period (0-10 s) from correct (blue) and error (red) trials at the left and right goals (WRS test, p = 0.0018 and < 0.001, respectively). (O-Q) Same as panels L to N but for the TRPC4 KD group (WRS test, p < 0.001 and < 0.001, for left and right trials, respectively in Q). Ctr, control; KD, knockdown.

Next, we investigated whether this affected the representations of the left and right goals at the population level using an SVM (Fig. 7D and E). This analysis was conducted using the activity of all cells from the 10 s goal period in each recording session with a minimum of 15 cells (n = 15 and 14 sessions in the control and KD groups, respectively). The left vs. right goal prediction accuracy and z-values of the SVM analysis were significantly lower in the TRPC4 KD group (Fig. 7D and E). While 80.0% of sessions had significantly higher prediction accuracy than chance level in control, only 42.8% were significantly higher than chance level in the KD group (Fig. 7E), suggesting that representations of left vs. right goals may have been compromised in the TRPC4 KD group.

We further investigated whether a gradual decline in left and right goal representation may be detected over time within the 10 s goal period owing to the lack of persistent activity in the TRPC4 KD group (Fig. 7F and G). We performed a similar SVM analysis at different time points starting from the pre-goal period (−3.5 s) using a sliding (1 s) 5 s window (time indicated in Fig. 7F and G is the center of the window). The prediction accuracy increased over time as the animal approached the goal area (−3.5 to +1.5 s, note that +1.5 still includes 3 s of pre-goal time). Importantly, the prediction accuracy and z-value between the control and KD groups differed starting at +4.5 s into the goal area (Fig. 7F and G), indicating a faster decline in goal representation in the KD group.

To test more directly the effect of TRPC4 KD on hippocampal spatial representations at the goal areas, we used Bayesian decoding of animal’s position based on the population activity (see Methods). In each trial, the actual position of the mouse (black trace) was compared with the decoded position (red trace) from 4 s before goal entry, when mice were in the stem and top of the T parts of the T-maze, until 8 s after the goal entry (Fig. 7H). While the position estimation was in general good in both control and TRPC4 KD group mice before the entry to the goal area, estimation errors increased in the TRPC4 KD group after the entry to the goal area (Fig. 7I). The estimation error between control and KD mice was not different before the entry to the goal area (−4 to 0 s; Fig. 7J) but was significantly larger in the KD group after the entry (0 to 8 s; Fig. 7K). This is in line with the data in the previous section where the spatial information was not different in the stem and the top of the T parts but was smaller in TRPC4 KD group in the goal areas. Together, these suggest the specific role of TRPC4 channels on the maintenance of spatial representation.

### Behavioral relevance

Finally, we asked whether the activity of the cells in the goal differed between correct and incorrect trials, using recording sessions with at least two error trials. The activity of the cells was first averaged over all trials, including both correct and error trials, to obtain combined sorting order. Subsequently, the activity profiles from correct and incorrect trials were separately averaged for each cell and sorted by the combined sort order (Fig. 7L). Persistent firing of the top 10% cells was significantly lower in the error trials compared to the correct trials in the control group for both the left and the right goals (Fig. 7M and N). Similar differences were observed in the KD group (Fig. 7O-Q). These results are consistent with the idea that hippocampal goal activity during the sample phases contributes to task performance^43,46^. Together, our data suggested that the intrinsic mechanism of persistent firing supports a stable spatial representation of goal locations, which is necessary for correct encoding of left vs right turns.

## Discussion

Despite empirical evidence^19,21,24,47^ and theoretical supports^26–28^, it remained unclear whether individual cells support persistent activity *in vivo*. We identified the TRPC4 channel as a molecular mechanism of intrinsic persistent firing in CA1 pyramidal cells^25^ and demonstrated that persistent firing was significantly reduced *in vivo* in a spatial working memory task by TRPC4 KD (Fig. 2-3). These results provide the first direct evidence that a single-cell mechanism supports persistent firing *in vivo* during a cognitive task. Further analyses revealed a specific effect of TRPC4 KD on persistently firing (Fig. 4) without disrupting signal-to-noise ratio in general. The behavioral outcome align with studies indicating the role of TRPC channels^48,49^ and manipulations that potentially disrupt persistent firing^32^ in working memory.

Classically, persistent firing during the delay period has been believed to hold working memory content. Abundant evidence supports this in the prefrontal cortex and hippocampus during the working memory delay period in humans and monkeys^5,8^ whereas data in rodents remains mixed^35,45,50^. In agreement with these rodent studies, our SVM analyses suggest the absence or very weak presence of working memory content in the delay phase (Fig. 5). An alternative theory posits that the hippocampus contributes to this type of working memory task by providing spatial information during the sample phase to help encoding, rather than maintaining working memory content^43,51,52^. We found that neurons with strong persistent firing had larger spatial information, and TRPC4 KD reduced spatial information specifically in places where persistent firing was observed (start and goal areas), suggesting that persistent firing mainly codes for spatial information (Fig. 6 and 7).

Hippocampal place cells are believed to be supported by path integration, which requires the maintenance and continuous updating of spatial representation^40^. Computational studies have suggested that similar attractor networks, such as continuous attractor networks, support working memory and path integration^37,38^. The continuous attractor model maintains spatial representation by persistent firing of a subset of neurons, “bump of activity”, through recurrent synaptic networks^38^. However, the neural networks of CA1, where place cells were most intensively studied, and MEC layer II, where grid cells were found, lack recurrent excitatory networks^53,54^. Additionally, these network models are sensitive to distractors, whereas experimental evidence suggests their robustness to distractors^55,56^.

Including the intrinsic cellular mechanisms of persistent firing in attractor networks has been shown to ameliorate these problems increasing stability against noise, reducing drift, and reducing sensitivity to parameters^28,29,57^. Upon empirically disrupting the intrinsic mechanism of persistent firing *in vivo*, we observed that spatial representation was compromised specifically at the start and goal areas. The intactness of spatial representation in other areas may indicate that attractor dynamics were still present, in agreement with the hybrid model. The gradual degradation of goal representation in the TRPC4 KD mice (Fig. 7) suggests a drift of attractor state due to the loss of persistent firing mechanism in individual cells as predicted from the loss of intrinsic properties in the hybrid model^28^.

An intact spatial representation outside the start and goal areas can be supported by external cue-based navigation^58^ that may not require information retention by persistent firing. Distal cues on the surrounding curtains and walls were visible to animals when they were in long corridors such as the stem and return arms, which was not the case in the start and goal areas. Alternatively, this could be attributed to the time required for persistent firing initiation, owing to the Ca2+-dependent biochemical processes involved in activating TRPC channels^20^. When the mice ran through the T-maze arms at a relatively high speed, the resulting short period of activation of each place cell may not have been long enough to fully activate the TRPC channels. Further studies are needed to clarify the exact mechanisms TRPC4 supports spatial representation in specific areas.

The prevailing view that neurons fire only when they receive sufficient input dictates current understanding of neural computation in the brain. Our findings challenge this view, revealing them as active contributors to information retention *in vivo*. This intrinsic persistent firing depends on the cholinergic system, which degrades in aging and Alzheimer’s disease^59,60^ causing cognitive impairments ^61,62^. Our findings thus potentially reshape our general understanding of brain computation and highlights novel therapeutic targets for cognitive disorders.

## Methods

### Animals

Twelve-week-old C57BL6/J male mice were obtained from a local animal facility. The mice were group-housed until the time of surgery and individually housed thereafter to avoid injury to sutures and/or implants. A 12-h light/dark cycle was used, and the mice underwent experiments during the light cycle. Mice housing room conditions were maintained at 21-24°C and 40-50% humidity. All experimental procedures were performed in accordance with animal research permission (TVA) obtained from the animal committee of Sachsen-Anhalt (Landesverwaltungsamt Referat Verbraucherschutz, Veterinärangelegenheiten, Az. 42502-2-1388 DZNE, 203.m-42502-2-1665 DZNE).

### TRPC4 channel knockdown

We developed a TRPC4 shRNA-expressing virus (TRPC4_AAV_U6_shRNA2_GFP) to knock down TRPC4 channels and a scramble virus (AAV_U6_scramble_GFP) for the control experiments. Oligos for TRPC4 were designed based on Puram and colleagues^63^ and were ordered from Sigma-Aldrich. The following oligonucleotides were used to clone siRNA against mTRPC4 into AAV_U6_GFP: mTRPC4_siRNA2: siRNA_TRPC4_Mouse/Human_2: Sense and antisense

5’-GATCC GGTCAGACTTGAACAGGCAA TTCAAGAGA TTGCCTGTTCAAGTCTGACC TTTTTTG - 3’

3’-G CCAGTCTGAACTTGTCCGTT AAGTTCTCT AAC-GGACAAGTTCAGACTGGAAAAAACTTAA-5’

Adeno-associated virus (AAV2) and U6 were used as the vectors and promoters, respectively. After purification, the titers were checked by qPCR (using TaqMan probes) and different dilutions. The titer for TRPC4_AAV_U6_shRNA2_GFP was 4.03*10^12^ and for AAV_U6_scramble_GFP was 3.56*10^12^. Finally, primary hippocampal cell cultures were infected with either no virus, scrambled virus, or TRPC4 KD virus, and a significant reduction in TRPC4 mRNA expression by the TRPC4 KD virus was confirmed by RT-qPCR (Fig. 1A).

### Surgery

C57BL/6JCrl male mice were weighed and anesthetized using a ketamine and xylazine cocktail (ketamine 20 mg/mL + xylazine 2.5 mg/mL, 0.1 mL/20 g). The head was fixed in the stereotactic device approximately 10 min after anesthetic injection. Head hair was completely removed, and the head was scrubbed using betadine (povidone-iodine) and ethanol 70%. A small incision was made using a scalpel, and the tissues were gently retracted. The skull position was checked horizontally. The Bregma point was used as reference, and the injection sites were marked according to coordinates from Paxinos atlas (AP: −2.1, ML±1.7, DV: 1.1 and AP: −2.7, ML±2.5, DV: 1.5). The marked points were drilled with a 0.7 mm drill tip. TRPC4_AAV_U6_shRNA2_GFP or AAV_U6_scramble_GFP viral particles were injected using a Hamilton neuro syringe (1701, 33 g needle) and stereotaxic syringe pump (CHEMYX, NANOJET) to control injection speed (100 nL/min) and volume (1 uL/site). The needle remained at the injection site for 5 min before retraction. Next, the skin was closed using veterinary surgery glue, and carprofen (5 mg/kg) was subcutaneously injected. Metamizole (200 mg/kg) was added to a 100 mL water bottle. The mice were allowed to recover for a few hours in the surgery room and then transferred to the animal room. They were checked daily (for three days) for recovery and pain management. For *in vivo* electrophysiological experiments, a microdrive with eight tetrodes was implanted on the right hemisphere and lateral to the virus injection site (AP: −2.1, ML±1.8, DV: 0.6). Four anchor screws and one ground screw are affixed to the skull. The implant was fixed using acrylic dental cement.

### *In vitro* neural recordings

The mice were deeply anesthetized with isoflurane, and the brains were quickly extracted after decapitation. Brains were placed in ice-cold cutting solution (110 mM) 110 Choline Cl, 7 mM MgCl2 (6H2O), 0.5 CaCl2, 2.5 KCl, 25 glucose, 1.2 NaH2PO4, 25 mM NaHCO3, 3 mM pyruvic acid, 11.5 ascorbic acid, and 100 mM D-mannitol). A Leica VT-1000 vibratome was used to obtain 350^M coronal slices. Brain slices were transferred to a holding chamber containing normal ACSF (nACSF) (126 mM) (126 mM NaCl, 1.2 mM NaH2PO4, 26 mM NaHCO3, 1.5 mM MgCl2, 1.6 mM CaCl2, 3 mM KCl, and 10 mM glucose). Slices were submerged for 30 min at 37 °C and then transferred to another holding chamber at room temperature for at least 30 min before recording.

Brain slices were transferred to the recording chamber and continuously superfused with nACSF (34.5 ± 1.5°C). The cells were visualized using an upright microscope (Olympus BX51WI) with a 40x water immersion objective and a Thorlabs CS895MU monochrome camera. GFP+ cells were located by exciting GFP with a 470 nm wavelength LED (CoolLED PE 4000) through a filter cube with excitation filter 469 ± 17.5 nm, dichroic 452-490 nm / 505-800 nm, and emission filter 525 ± 19.5 nm. Whole-cell configuration was achieved using 3-5 MΩ patch pipettes filled with intracellular solution (in mM: 120 K-gluconate, 10 HEPES, 0.2 EGTA, 20 KCl, 2 MgCl2, 7 PhCreat di(tris), 4 Na2ATP, and 0.3 Tris-GTP (pH adjusted to 7.3 with KOH). For labeling purposes, 0.1% biocytin was added to the intracellular solution. Access resistance was monitored and compensated for several times during recording. The liquid junction potential was not corrected. Electrophysiological signals were amplified using a Multiclamp 700A amplifier and digitized using an Axon Digidata 1550 B. Current clamp recordings were routinely filtered at 10 KHz and sampled at 20 KHz using pClamp 11 software. After recording, the slices were stored at 4°C in a phosphate-buffered saline (PBS) solution with 4% paraformaldehyde (PFA).

### Behavioral task

The spatial working memory task in the T-maze was started four weeks after virus injection. Before this test, the mice were handled for three days to habituate to the experimenter. Mice were food-deprived at the start of handling to motivate them for the reward, and reward pellets were introduced to them before starting the main experiment. Animal weight was maintained between 85-90% of body weight before food deprivation. Second, habituation to the T-maze (OHARA Ltd., Japan) was performed for three days.

Each animal was tested for 10 days, with each day constituting 10 trials (10 sample +10 choice phases). Each trial started with a sample trial (forced trial), in which the T-maze software assigned the left or right door (at the T-junction) to open randomly. First, the start door was opened, and the mouse moved from the start area to the stem, open arm (top of the T-maze), and goal area where it received the reward. Next, the animal returned to the start area using the return arm and waited for 30 s (delay period). After the delay, the choice trial began; this time, both the left and right arms were open, and the mouse chose the arm opposite the sample trial. If the trial was successful, the animal received a pellet at the goal location. After collecting the reward, the animal moved back to the start area, where a new trial began after 45 s (inter-trial interval, ITI). After each experiment, the maze floor was cleaned with 10% ethanol, and the walls were cleaned with water. Behavioral data were analyzed using MATLAB (MathWorks) and SPSS (GEE).

### *In vivo* neural recordings

Tetrodes were made using 12.5 μm Tungsten wire (California Fine Wire Co.) and grouped in a single bundle with eight tetrodes (32 channels) inside a microdrive (AXONA). The tetrodes were gold-plated using nanoZ (White Matter LLC, USA) in gold solution to reduce the impedance to 150 kQ.

Screening was conducted 7-10 days after surgery and after handling the animals to place the tetrodes in the pyramidal layer of CA1. Screening was performed in an open-field arena (30 x 30 cm, shielded and grounded). The presence of place cells, autocorrelation, firing frequency, and waveform characteristics were the criteria used to identify the pyramidal layer. Once the tetrodes were in a position or very close to CA1, the behavioral experiment in the T-maze began.

Single-unit activity and local field potentials (LFP) were recorded using an in vivo electrophysiology setup (Axona Ltd.). The data were recorded in the raw mode at a sampling rate of 48 kHz. To obtain single-unit spike data, raw data were referenced and band-passed (400-8000 Hz) using custom scripts written in MATLAB (MathWorks), and spikes were detected using TINT (Axona Ltd.). The detected spikes were clustered using KlustaKwik^64^ for automatic clustering, and manual curation of the clusters was performed using the Klusters software (Lynn Hazan). LFP data with a 4800 Hz sampling rate were created using TINT (Axona Ltd.). The position of the mouse was tracked at 50 Hz using a video camera that recorded the position of a light-emitting diode attached to the mouse head.

### Histology

Mice were anesthetized with a ketamine and xylazine cocktail and transcardially perfused with cold PBS (4°C, 15 mL) and then cold paraformaldehyde 4% (PFA, PH∼7.4, 50 mL). Brains were removed and post-fixed in PFA 4% at 4°C for 24 h. After applying the cryoprotective protocol (one day in sucrose 10%, three days in sucrose 30%), samples were embedded in cubic molds filled with Optimal Cutting Temperature compound (O.C.T.) and then snap frozen. Next, the samples were transferred to the −80°C freezer for storage. The fixed samples were sectioned using a cryostat. The slices were then covered and mounted using a mounting medium. Images were obtained with a confocal microscope (Zeiss, AxioImager M2) and fluorescence microscope (Keyence, BZ-X710) using 2.5x, 5x, and 40x objectives to control for virus expression and tetrode traces (Supplemental Fig. 1).

### *In vitro* data analysis

The resting membrane potential was measured by averaging the membrane potential for 1 min after seal opening without current injection. Input resistance was calculated using Ohm’s law, measuring the voltage deflection in response to a negative current pulse of −50 pA at −65 mV. Intrinsic excitability was evaluated by counting the number of action potentials during the positive step-wise current injections (1 s), from −300 to 300 pA in 50 pA jumps (IV protocol). The adaptation index was calculated by dividing the instantaneous frequency of the last pair of action potentials by the instantaneous frequency of the second pair of action potentials in the first sweep, with at least 10 action potentials in the IV protocol. *In vitro* persistent firing was induced by a 2 s 100 pA current injection at a voltage level of 5­3 mV below the action potential threshold. The strength of persistent firing was evaluated by calculating the average firing frequency and depolarization in the first 10 s post-stimulus. The cells were classified as having long (> 30 s), short (< 30 s), or no persistent firing, based on the duration of persistent firing observed (measured from the offset of the stimulus).

### *In vivo* data analysis

The data were analyzed using custom-written MATLAB scripts and the CMBHome toolbox developed at the Center for Memory and Brain, Boston University.

#### Temporal binning and Z-score firing rate

The time-binned firing rate of each cell was prepared using a 0.5 s bin and smoothed with a 2.5 s Gaussian window to analyze *in vivo* persistent firing. The Z-scored firing rate was calculated from the time-binned firing rate data using the following equation:

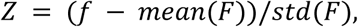

where *f* and F are the binned firing rates from the period of interest and baseline period, respectively.

#### Cell type classification based on activity change

Neurons were classified into activated (Act), suppressed (Sup), and non-modulated (NM) cells, according to the delay period activity^65^ (Fig. 2E). The binned firing rate at baseline (15-5 s before the delay) was compared to the activity during the delay (0-25 s) using the Wilcoxon rank-sum test (WRS). Cells were considered to be modulated if the test result was significant (p < 0.01). The z-value obtained during the test (WRS z-value) was used to classify the modulated cells into the Act (WRS z-value > 2.58) and Sup (WRS z-value < −2.58) groups. The remaining cells were classified as NM. The WRS z-values were used to plot a histogram, as shown in Fig. 2E.

#### Temporal field

The peak z-scored firing rate during the delay period was determined to define the temporal field for each cell. The temporal field duration was defined as the time when the z-scored firing rate was continuously above 10% of the peak value (Fig. 2I). In Fig. 3, the temporal field duration is defined in the same way, using the raw firing rate instead of the z-scored firing rate.

#### SVM analysis

A population vector was built for each trial of each session using neuronal activity during the delay period (0-25 s) or from the goal area (0-10 s). Single-unit spike activity was temporarily binned (0.5 s bin) for left vs. right and ITI vs. delay comparisons. A temporal bin size of 0.02 s was used for the left vs. right goal comparison with a 5 s sliding window (Fig. 7F and G). Each trial was then labeled according to the type of test: ITI vs. delay or left vs. right (retrospective or prospective). Only sessions with at least 15 units were included in the analysis.

Binary SVM comparisons were performed for each session with the fitcsvm function (MATLAB) using a linear kernel and two-fold cross-validation of the input data to evaluate model performance. The model was run 30 times, and the obtained balanced accuracies were averaged. To evaluate the significance of the prediction, the class labels were scrambled, and the SVM model was run for the non-scrambled data a minimum of 30 times (sessions where label permutation did not generate at least 30 different combinations were not included in the analysis). Moreover, a random accuracy distribution was built and then used to determine the z-scored balance accuracy. Sessions with z-score > 1.96 were considered to have above-chance accuracy.

#### Spatial information and place cells

Spatial information (SI) is defined as the extent to which the firing of a cell can be used to predict the position of the animal. The SI was calculated using the formula of Skaggs et al. (1992)^66^:

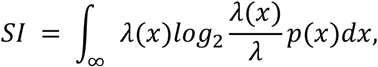

where SI is the information rate in bits/s, x is the spatial location of the animal, A(x) is the mean frequency at location x, p(x) is the probability of finding the animal at a given location in the maze, and *A* is the overall firing rate of the cell.

Spatially modulated cells were selected by building a randomized distribution of spatial information values obtained by circularly shifting the spike timing and position vectors for at least 20 s 100 times. Cells with spatial information above the top 95% of the random distribution were considered spatially modulated cells (place cells).

The spatial rate maps represent the average firing rate of a cell at a given location, and were obtained by dividing the number of spikes at a location by the total time that the animal spent at that location (occupancy). Rate maps were calculated using 1×1 cm bins, and the data were smoothed using a two-dimensional convolution with a Gaussian kernel with a 2.5 cm standard deviation. Only bins with speed greater than 2.5 cm/s were considered in the calculation.

Place field centers were detected as follows. First, pixels of a rate map with a high firing rate (> 4 SD) were extracted. Extracted rate map was processed with the regionprops function (MATLAB) to detect and characterize the individual fields and the area and centroid parameters were obtained. To avoid detecting noise, fields smaller than 12 or larger than 650 cm^2^ were discarded, and firing field with the highest coefficient of variation was considered the main firing field of a given neuron. The coefficient of variation was calculated as follows:

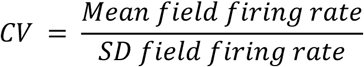

#### Theta amplitudes

Theta components were extracted by bandpass filtering of the LFP data (6-10 Hz). The theta amplitude was calculated using the absolute value of the Hilbert transform of the filtered signal.

#### Putative cell type classification

Putative pyramidal neurons and interneurons were identified based on the average firing rate and spike duration. Cells with an average firing rate < 8 Hz and a peak-to-peak spike duration > 0.255 ms were classified as putative pyramidal cells, whereas those with an average firing rate > 8 Hz and a peak-to-peak spike duration < 0.255 ms were classified as putative interneurons^67,68^.

#### Trial selection for goal analysis

The duration of time animals stayed at the goal areas varied greatly from trial to trial, ranging from 1.6 to 410.0 s (median duration: 47.74 and 47.73 s for control and TRPC4 KD groups, respectively; WRS test, p = 0.99), because the task did not restrict the time animals spend at the goal areas. In addition, the animals returned to the top arm (top of T) in some trials. Therefore, we limited our goal analysis to trials in which the animals continuously spent at least 10 s within the goal area during the sample phase. When multiple such periods were found in a trial, we analyzed the first incident. The baseline period to build the z-scored firing rate was the last 20 s of the ITI period before goal entry. This ensured that goal activity was not included in the baseline period. In Fig. 7A, H, and K, the black vertical gap separates the first 10 s, which is referenced to the end of the ITI period, from the rest, which is referenced to the start of goal entry. The trials in each recording session were first divided into left and right sample goal trials to compare correct and incorrect trial activities. Subsequently, we checked whether at least one correct or incorrect trial was present for each goal location. Sessions that did not meet these criteria were excluded from the analysis.

#### Bayesian position decoding

For Bayesian position decoding, we included sessions with at least 30 simultaneously recorded CA1 units. In each session, position decoding was performed on trials in which the mouse stayed at the goal location for longer than 10 s because our focus was on the maintenance of spatial representation over time. In each trial, position decoding was performed using a 12-s window spanning 4 s before to 8 s after goal entry (goal window). To build spatial tuning curves for later decoding, two-dimensional rate maps with a 1.7 cm spatial bin size were calculated for all cells using the spike data from the complement of the goal window within the session (i.e., all spike data of the session excluding the goal window from corresponding trial). Data used for decoding in each goal window consisted of spike counts from all cells binned into non-overlapping 200 ms time bins (Δt = 0.2 s), which were smoothed across time using a Gaussian kernel (6-bin window).

Decoding assumed independent Poisson spiking across units, such that for each spatial bin (state) *s* with expected rate X_i_(*s*) from the rate map, the log-likelihood of observing counts *r*_i_ in a time bin was computed as £_i_[*r*_i_log X_i_(*s*) - X_i_(*s*)A*t*]; rate-map values were floored at 10^-6^ Hz to ensure numerical stability. A uniform prior over spatial states was used, and the posterior over all spatial bins was obtained by combining log-likelihood and log-prior and normalizing each time bin.

Decoded position was taken as the maximum a posteriori (MAP) spatial bin and converted to x-y coordinates using bin centers; posterior marginals over x and y were additionally computed by summing the 2D posterior over the orthogonal dimension. Ground-truth x(t) and y(t) at each decoding time bin midpoint was obtained by linear interpolation of the tracked position and discretized into a spatial bin center coodinate. Decoding accuracy was quantified as the Euclidean distance between decoded and ground-truth position at each time bin.

### Statistical analysis

The data were checked for distribution and homogeneity before statistical analysis. Comparisons between groups were performed using the Wilcoxon rank-sum and Kruskal-Wallis tests. A t-test or analysis of variance (ANOVA) followed by Tukey’s post-hoc test was used for normally distributed data. Multiple-comparison corrections were used where relevant. Categorical variables (group ratios) were compared using the chi-squared test or Fisher’s exact test if any group had fewer than five items. Boxplots represent the median and upper and lower quartiles. Significance was expressed as follows: * p < 0.05, ** p < 0.01, and *** p < 0.001.

## Data availability

Data used in the current study will be available on request to the corresponding authors.

## Code availability

All source codes used in the current study will be available on request to the corresponding authors.

## Acknowledgments

We would like to thank K. Jeffery, S. Leutgeb, J. Leutgeb, M. Brandon, and K. Allen for their help in establishing the *in vivo* electrophysiological recordings. We thank L. Leichert and O. Stork for their assistance and technical support, and M. Freichel for sharing the TRPC KO mice with us. We thank M. Hasselmo, B. McNaughton, T. McHugh and S. Remy for their valuable comments on this manuscript. This work was supported by the Federal State of Saxony-Anhalt and the European Structural and Investment Funds (ESF, 2014-2020, project number ZS/2016/08/80645), the Deutsche Forschungsgemeinschaft projects YO177/4-1, YO177/4-3, and YO177/7-1 (to M.Y.), and the German Center for Neurodegenerative Diseases (to A.D.).

## Author Contributions

Conceptualization and experimental design, M.Y. and M.S.; analysis, A.R., M.Y., and B.S.M.; investigation, B.S.M.; writing, M.Y. and A. R.; methodology, A.D. and R.K.; software, M.Y. and F.T.; funding acquisition, M.Y.; supervision, M.Y. and M.S.

## Competing interests

The authors declare no competing interests.

## Supplemental Information

**Supplemental Figure 1.**
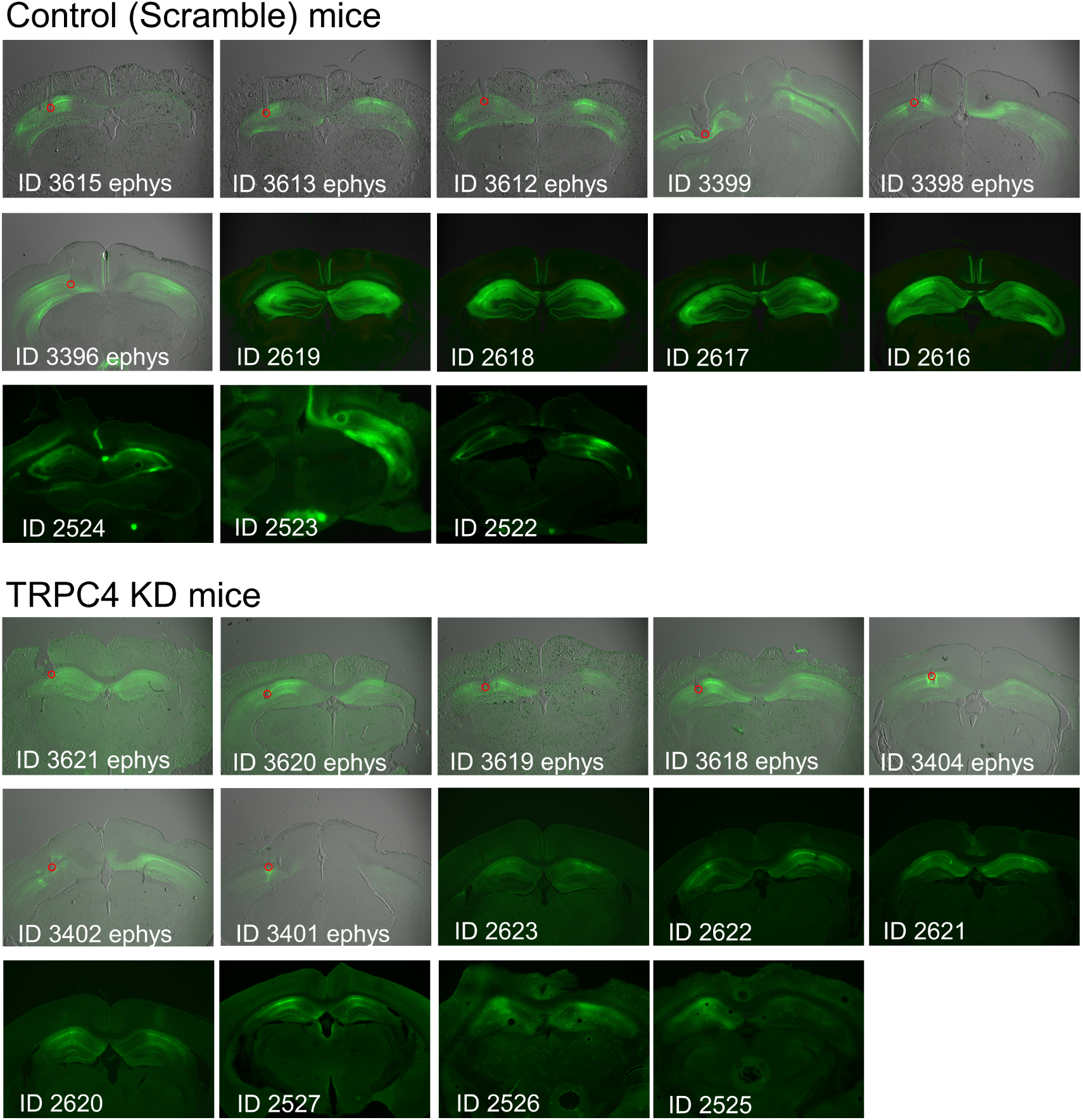
Virus expression and tetrode track GFP expression and track locations of each animal for both the control and TRPC4 KD groups. Animal IDs followed by “ephys” indicate that those mice were used for electrophysiological recording. The tips of tracks are highlighted with a red circle. Other animals were only used for the behavioral test.

**Supplemental Figure 2.**
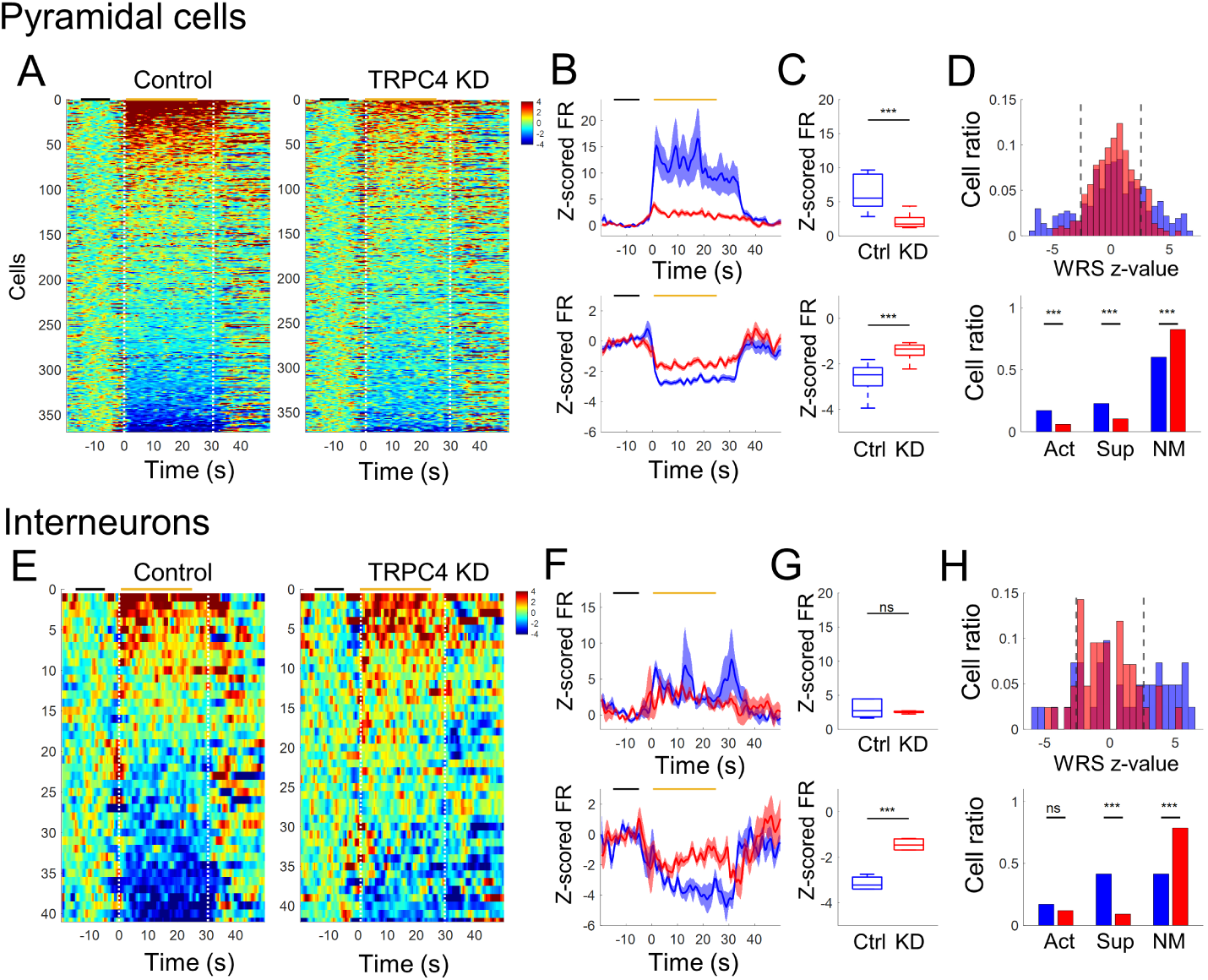
Activity of putative pyramidal and interneurons (A) Z-scored firing rate of putative-pyramidal cells sorted by the delay period activity. Black and orange horizontal lines at the top of each panel indicate baseline period and period for cell sorting, respectively. (B) Averaged z-scored firing rate (FR) of top (upper panel) and bottom (lower panel) 10% cells in A. (C) Comparison of z-scored firing rate of the top (upper panel; Wilcoxon rank sum Test, p < 0.001) and bottom (lower panel; Wilcoxon rank sum test, p < 0.001) 10 % cells in A during the delay period (0-25 s). (D) Classification of activated (Act), suppressed (Sup) and non-modulated (NM) cells using Mann-Whitney U (MWU) test. The upper panel shows the distribution of Wilcoxon rank-sum z-values and the lower panel shows the ratio of cells in each category (WRS test, p < 0.001 for all groups). (E to H) Same as A to D but for putative-interneurons. Statistical tests in G: WRS test, p = 0.6905 in the upper panel, and p < 0.001 in the lower panel.

**Supplemental Figure 3.**
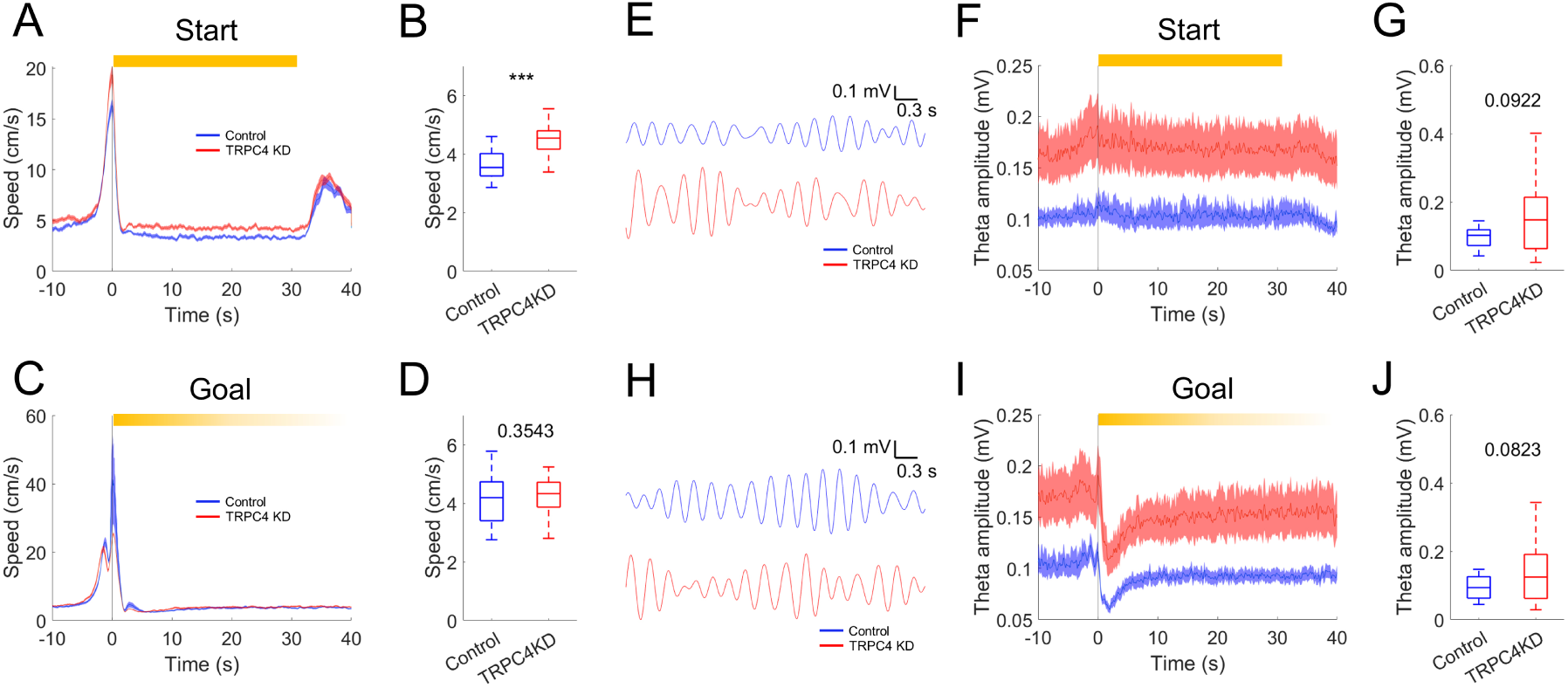
Running speed and theta amplitude at the start and goal areas (A) Average running speed of control and TRPC4 KD animals at the start area. Time is relative to the entry to the start area. This figure includes both the sample (ITI) and choice (delay) phases. (B) Comparison of the speed in the start area (Wilcoxon rank sum test, p < 0.001). (C) Same as A but for the goal area. Note that there was no set duration during which animals stayed at the goal area. While we show 40 s of goal speed data, animals were already out of the goal areas in some trials. (D) Comparison of the speed in the goal area (Wilcoxon rank sum test, p = 0.35). For this comparison, the speed of each session was calculated using only the time the animal stayed in the goal area. (E) Examples of filtered theta oscillation at the start area. (F) Theta amplitudes of control and TRPC4 KD animals at the start area. (G) Comparison of the averaged theta amplitudes in the start area (Wilcoxon rank sum test, p = 0.092). (H) Examples of filtered theta oscillation at the goal area. (I) Theta amplitudes of control and TRPC4 KD animals at the goal area. As in C, animals were out of the goal areas in some trials before the end of the trace. (J) Comparison of the theta amplitudes in the goal area (Wilcoxon rank sum test p = 0.082). As in D, the theta amplitude of each session was calculated using only the time the animal stayed in the goal area.

## Notes

### Competing Interest Statement

The authors have declared no competing interest.

### Summary of Updates

new title, abstract, and introduction new analysis on position decoding

